# Adaptive evolution of nontransitive fitness in yeast

**DOI:** 10.1101/700302

**Authors:** Sean W. Buskirk, Alecia B. Rokes, Gregory I. Lang

**Affiliations:** Department of Biological Sciences, Lehigh University, Bethlehem PA 18015; Department of Biology, West Chester University, West Chester PA 19383

## Abstract

Nontransitivity – commonly illustrated by the rock-paper-scissors game – is well documented among extant species as a contributor to biodiversity. However, it is unclear if nontransitive interactions also arise by way of genealogical succession, and if so, through what mechanisms. Here we identify a nontransitive evolutionary sequence in the context of yeast experimental evolution in which a 1,000-generation evolved clone outcompetes a recent ancestor but loses in direct competition with a distant ancestor. We show that nontransitivity arises due to the combined forces of adaptation in the yeast nuclear genome and the stepwise deterioration of an intracellular virus. We show that, given the initial conditions of the experiment, this outcome likely to arise: nearly half of all populations experience multilevel selection, fixing adaptive mutations in both the nuclear and viral genomes. In contrast to conventional views of virus-host coevolution, we find no evidence that viral mutations (including loss of the virus) increase the fitness of the host. Instead, the evolutionary success of evolved viral variants results from their selective advantage over viral competitors within the context of individual cells. Our results provide the first mechanistic case-study of the adaptive evolution of nontransitivity, in which a series of adaptive replacements produce organisms that are less fit when compared to a distant genealogical ancestor.

## Introduction

Adaptive evolution is a process in which selective events result in the replacement of less-fit genotypes with a more fit ones. Intuitively, a series of selective events, each improving fitness relative to the immediate predecessor, should translate into a cumulative increase in fitness relative to the ancestral state. However, whether or not this is borne out over long evolutionary time scales has long been the subject of debate (Ruse 1993, Dawkins 1997, Gould 1997, Shanahan 2000). The failure to identify broad patterns of progress over evolutionary time scales—despite clear evidence of selection acting over successive short time intervals—is what Gould referred to as “the paradox of the first tier (Gould 1985).” This paradox implies that evolution exhibits nontransitivity, a property that is best illustrated by the Penrose staircase and the Rock-Paper-Scissors game. The Penrose staircase is a visual illusion of ascending sets of stairs that form a continuous loop such that—although each step appears higher than the last—no upward movement is realized. In the Rock-Paper-Scissors game each two-way interaction has a clear winner (paper beats rock, scissors beats paper, and rock beats scissors), yet due to the nontransitivity of these two-way interactions, no clear hierarchy exists among the three.

In ecology, nontransitive interactions among extant species are well-documented as contributors to biological diversity and community structure (Kerr et al. 2002, Károlyi et al. 2005, Laird and Schamp 2006, Reichenbach et al. 2007, Menezes et al. 2019) and arise by way of resource (Sinervo and Lively 1996, Precoda et al. 2017) or interference competition (Kirkup and Riley 2004). First put forward in the 1970s (Gilpin 1975, Jackson and Buss 1975, May and Leonard 1975, Petraitis 1979), the importance of nontransitivity in ecology has garnered extensive theoretical and experimental consideration over the last half century (e.g. Sinervo and Lively 1996, Kerr et al. 2002, Allesina and Levine 2011, Rojas-Echenique and Allesina 2011, Soliveres et al. 2015, Liao et al. 2019).

What is unknown is whether nontransitive interactions arise for direct descendants along a line of genealogical succession. This is the crux of Gould’s paradox and has broad implications for our understanding of evolutionary processes. For instance, if an evolved genotype is found to be less fit in comparison to a distant ancestor, the adaptive landscape upon which the population is evolving may not contain true fitness maxima (Barrick and Lenski 2013, Van den Bergh et al. 2018) and, more broadly, directionality and progress may be illusory (Gould 1996). Testing the hypothesis that nontransitive interactions arise along lines of genealogical descent, however, is not possible in natural populations because it requires our ability to directly compete an organism against its immediate predecessor as well as against its extinct genealogical ancestors. Fortunately, laboratory experimental evolution, in which populations are preserved as a “frozen fossil record,” affords us with the unique opportunity to test for nontransitivity along a genealogical lineage by directly competing a given genotype against the extant as well as the extinct.

An early study of laboratory evolution of yeast in glucose-limited chemostats appeared to demonstrate that nontransitive interactions arise along a line of genealogical descent (Paquin and Adams 1983). However, the specific events that led to nontransitivity in this case are unknown, and it is likely the case that the authors were measuring interactions between contemporaneous lineages in a population, rather than individuals along a direct line of genealogical descent, as they report (see Discussion). Indeed, adaptive diversification is common in experimental evolution due to spatial structuring (Rainey and Travisano 1998, Frenkel et al. 2015) and metabolic diversification (Helling et al. 1987, Turner et al. 1996, Spencer et al. 2008, Plucain et al. 2014), and is typically maintained by negative frequency-dependent selection, in which rare genotypes are favored. Collectively this work reinforces theory and observational evidence on the power of ecological nontransitivity as a driver and maintainer of diversity but is silent as to whether genealogical succession can also be nontransitive.

Here we determine the sequence of events leading to the evolution of nontransitivity in a single yeast population during a 1,000-generation evolution experiment. We show that nontransitivity arises through multilevel selection involving both the yeast nuclear genome and the population of a vertically-transmitted virus. Many fungi, including the yeast *Saccharomyces cerevisiae* are host to non-infectious, double-stranded RNA “killer” viruses (Wickner 1976, Schmitt and Breinig 2002, Schmitt and Breinig 2006, Rowley 2017). Killer viruses produce a toxin that kills non-killer containing yeasts. The K1 toxin gene contains four subunits (*δ, α, γ, β*), which are post-translationally processed and glycosylated to produce an active two-subunit (*α, β*) secreted toxin (Bostian et al. 1983). Immunity to the toxin is conferred by the pre-processed version of the toxin, thus requiring cells to maintain the virus for protection. We show that nontransitivity arises due to multilevel selection: adaptation in the yeast nuclear genome and the simultaneous stepwise deterioration of the killer virus. By expanding our study of host-virus genome evolution to over 100 additional yeast populations, we find that multilevel selection, and thus the potential for the evolution of nontransitive interactions, is a common occurrence given the conditions of our evolution experiment.

## Results

### Evolution of nontransivity along a line of genealogical descent

Previously we evolved ∼600 haploid populations of yeast asexually for 1,000 generations in rich glucose medium (Lang et al. 2011). We characterized extensively the nuclear basis of adaptation for a subset of these populations through whole-genome whole-population time-course sequencing (Lang et al. 2013) and/or fitness quantification of individual mutations (Buskirk et al. 2017).

For one population (BYS1-D08) we were surprised to observe that a 1,000-generation clone lost in direct competition with a fluorescently-labeled version of the ancestor. To test the hypothesis that a nontransitive interaction arose during the adaptive evolution of this population, we isolated individual clones from three timepoints (Fig. 1A). We define these as Early (Generation 0), Intermediate (Generation 335, chosen to coincide with the first clonal replacement, which fixes first three nuclear mutations), and Late (Generation 1,000, following three additional clonal replacements, which fix ten nuclear mutations). We performed pairwise competition experiments between the Early, Intermediate, and Late clones at multiple starting frequencies. We find that the Intermediate clone is 3.8% more fit relative to the Early clone and that the Late clone is 1.2% more fit relative to the Intermediate clone (Fig. 1B, left panel). The expectation, assuming additivity, is that the Late clone will be more fit than the Early clone, by roughly 5.0%. Surprisingly, we find that the Late clone is less fit than expected, to the extent that it often loses in pairwise competition with the Early clone (Fig. 1B, left panel). Furthermore, the interaction between the Early and Late clones exhibits positive frequency-dependent selection, thus creating a bistable system where the fitness disadvantage of the Late clone can be overcome if it starts above a certain frequency relative to the Early clone (Fig. S1).

**Fig. 1.**
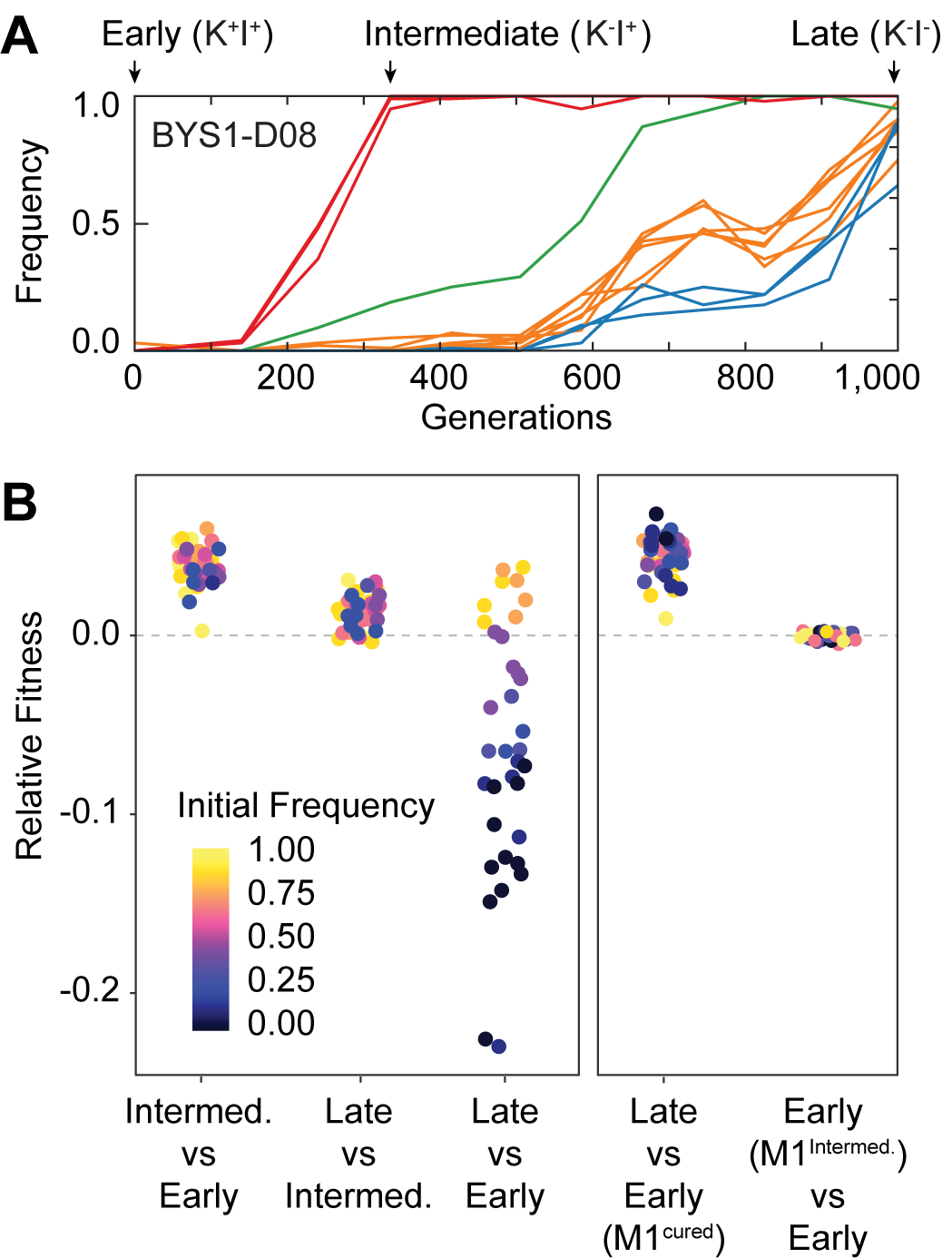
Nontransitivity and positive frequency dependence arise along an evolutionary lineage. A) Sequence evolution (from Lang *et al*. 2013) shows that population BYS1-D08 underwent four clonal replacements over 1,000 generations. Mutations in the population that went extinct are not shown. The four selective sweeps are color-coded: red, mutations in *yur1, rxt2*, and an intergenic mutation; green, a single intergenic mutation; orange, mutations in *mpt5, gcn2, iml2, ste4, mud1*, and an intergenic mutation; blue, three intergenic mutations. The Intermediate clone, isolated at Gen. 335 does not produce, but is resistant to, the killer toxin (K-I+). The Late clone, isolated at Generation 1,000 does not produce, and is sensitive to, the killer toxin (K-I-). B) Competition experiments demonstrate nontransitivity and positive frequency-dependent selection. Left: Relative fitness of Early (Gen. 0), Intermediate Gen. 335), and Late (Gen. 1,000) clones. Right: Relative fitness of the Early clone without ancestral virus or with viral variant from the Intermediate clone. Fitness and starting frequency correspond to the later clone relative to earlier clone during pairwise competitions.

### Evolution of nontransitivity is associated with changes to the killer virus

Positive frequency-dependent selection is rare in experimental evolution and can only arise throught a few known mechanisms. It has been observed previously in yeast that harbor killer viruses (Greig and Travisano 2008), which are dsRNA viruses that encode toxin/immunity systems. Using a well-described halo assay (Woods and Bevan 1968), we find that the ancestral strain of our evolved populations exhibits the phenotype expected of yeast that harbor the killer virus: it inhibits growth of a nearby sensitive strain and resists killing by a known killer strain (Fig. S2). By RT-PCR and sequencing we find that our ancestral strain, which was derived from the common lab strain W303-1a, contains the M1-type killer virus (encoding the K1-type killer toxin) with only minor differences from previously sequenced strains (Fig. S3). In the ancestor we also detect the L-A helper virus, which supplies the RNA-dependent RNA polymerase and capsid protein necessary for the killer virus, a satellite virus, to complete its life cycle (Ribas and Wickner 1992).

We asked if the observed nontransitivity in the BYS1-D08 lineage could be explained by evolution of the killer phenotype. Phenotyping of the isolated clones revealed that the Intermediate clone no longer exhibits killing ability (K^-^I^+^) and that the Late clone possesses neither killing ability nor immunity (K^-^I^-^, Fig. 1A, Fig. S4). Killer toxin has been shown to impart frequency-dependent selection in structured environments (Greig and Travisano 2008) and we hypothesized that a stepwise loss of the killer phenotypes was responsible for the frequency-dependent and nontransitive interaction between Early and Late clones. To determine if the presence of the killer virus in the early clone is necessary for the evolution of nontransitivity, we cured the Early clone and found that the Late clone was 4.3% more fit than the killer-cured Early clone, with no correlation between fitness and frequency (Fig. 1B, right panel) showing that the presence of killer virus in the Early clone is necessary for frequency-dependence and nontransitivity. To determine if viral evolution alone is sufficient to account for the observed fitness gains in nontransitive interactions, we transferred the killer virus from the Intermediate clone to the cured Early clone and assayed fitness relative to the Early clone (Fig. 1B, right panel). We find that the evolved killer virus confers no significant effect on host fitness. Therefore, changes to the killer virus alone are not sufficient to account for the adaptive evolution of nontransitivity in this population, which must involve changes to both the host and viral genomes.

### Changes to killer-associated phenotypes are common under our experimental conditions

To determine the extent of killer phenotype evolution across all populations, we assayed the killer phenotype of 142 populations that were founded by a single ancestor and propagated at the same bottleneck size as BYS1-D08 (Lang et al. 2011). We find that approximately half of all populations exhibit a loss or weakening of killing ability by Generation 1,000, with ∼10% of populations exhibit neither killing ability nor immunity (Fig. 2A, Fig. 2B). Of note, we did not observe loss of immunity without loss of killing ability, an increase in killing ability or immunity, or reappearance of killing ability or immunity once it was lost from a population (Fig. S5), apart from the noise associated with scoring of population-level phenotypes. Several populations (i.e. BYS2-B09 and BYS2-B12) lost both killing ability and immunity simultaneously, suggesting that a single event can cause the loss of both the killer phenotypes.

**Fig. 2.**
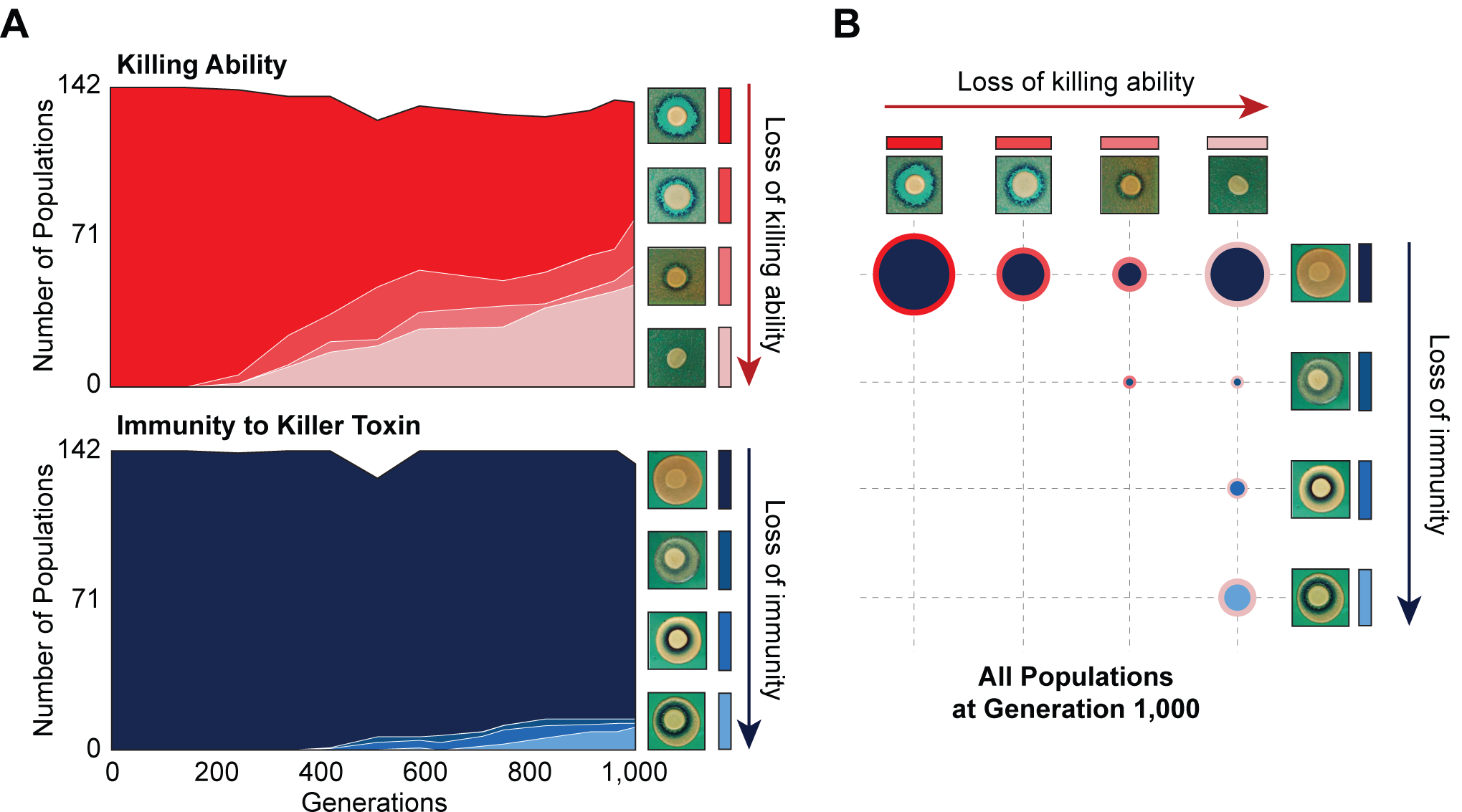
Changes in killer-associated phenotypes in the 142 populations that were founded by a single ancestor and propagated at the same bottleneck size as BYS1-D08 (Lang *et al*. 2011). A) Loss of killing ability (top) and immunity (bottom) from evolving yeast populations over time. Killer phenotypes were monitored by halo assay (examples shown on right). B) Breakdown of killer phenotypes for all populations at Generation 1,000. Data point size corresponds to number of populations. Border and fill color indicate killing ability and immunity phenotypes, respectively, as in panel A.

Mutations in nuclear genes can affect killer-assiciated phenotypes. The primary receptors of the K1 killer toxin are β-glucans in the yeast cell wall (Pieczynska et al. 2013). We observe a statistical enrichment of mutations in genes involved in β-glucan biosynthesis (6-fold Gene Ontology (GO) Biological Process enrichment, *P*<0.0001). Furthermore, of the 714 protein-coding mutations dispersed across 548 genes, 40 occur within 11 of the 36 genes (identified by (Pagé et al. 2003)) that, when deleted, confer a high level of resistance to the K1 toxin (χ^2^=18.4, *df*=1, *P*=1.8×10^−5^). Nevertheless, the presence of mutations in nuclear genes that have been associated with high levels of resistance is not sufficient to account for the loss of killing ability (χ^2^=1.037, *df*=1, *P*=0.309) or immunity (χ^2^=0.103, *df*=1, *P*=0.748).

### Standing genetic variation and *de novo* mutations drive phenotypic change

We sequenced viral genomes from a subset of yeast populations (n=67) at Generation 1,000, 57 of which change killer phenotype and 10 control populations that retained the ancestral killer phenotypes. Viral genomes isolated from populations that lost killing ability possess 1-3 mutations in the M1 coding sequence – most being missense variants (Fig. 3A). In contrast, only a single mutation, synonymous nonetheless, was detected in M1 across the 10 control populations that retained the killer phenotype (χ^2^=59.3, *df*=1, *P*=1.4×10^−13^). The correlation between the presence of mutations in the viral genome and the loss of killing ability is strong evidence that viral mutations are responsible for the changes in killer phenotypes. We estimate that by Generation 1,000 half of all populations have fixed viral variants that alter killer phenotypes (for comparison, *IRA1*, the most common nuclear target, fixed in ∼25% of populations over the same time period).

**Fig. 3.**
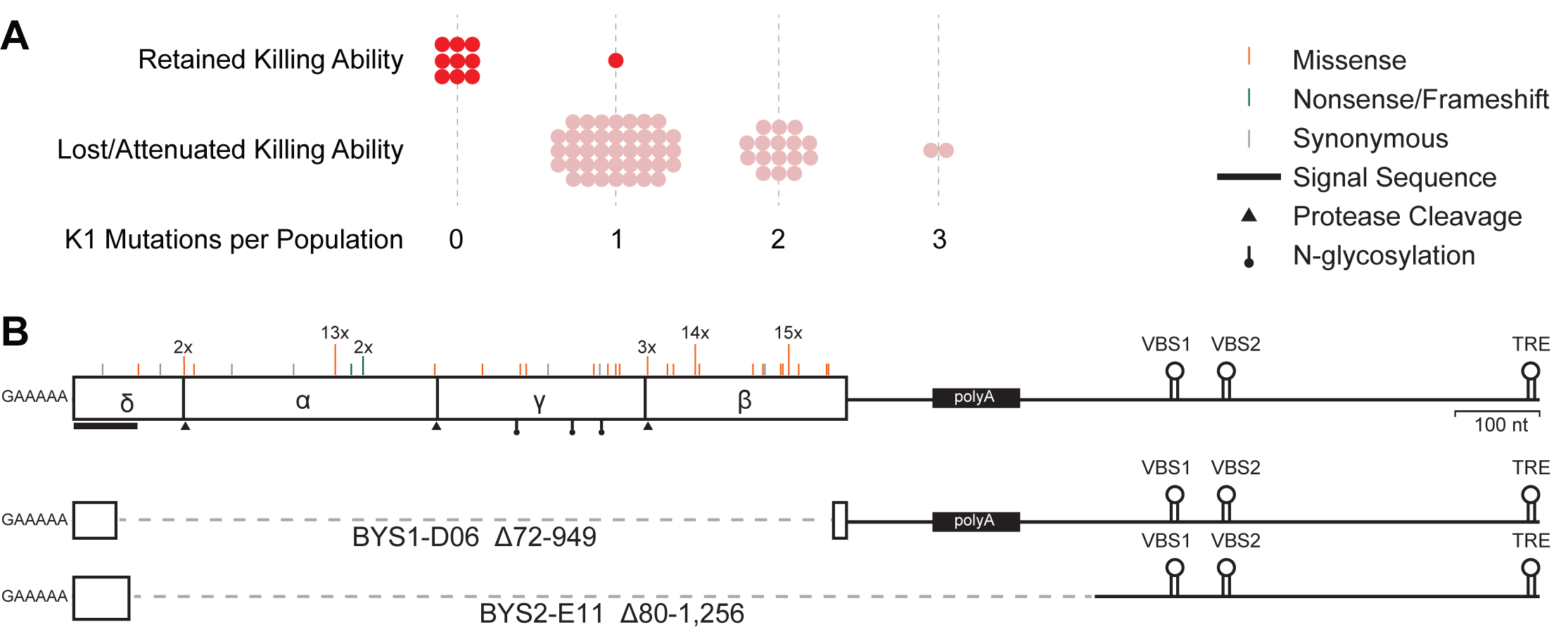
Loss of killer phenotype correlates with presence of mutations in the K1 toxin gene. A) Number of mutations in the K1 gene in yeast populations that retain or lose killing ability. Each data point represents a single yeast population. B) Observed spectrum of point mutations across the K1 toxin in 67 evolved yeast populations. Mutations were detected in a single population unless otherwise noted. Large internal deletion variants from two yeast populations (BYS1-D06 and BYS2-E11). The deletions span the region indicated by the dashed gray line. VBS: viral binding site. TRE: terminal recognition element.

Of the 57 populations that lost killing ability, 42 fixed one of three single nucleotide polymorphisms, resulting in amino acid substitutions D106G, D253N, and I292M and observed 13, 14, and 15 times, respectively (Table S1). Given their prevalence, these polymorphisms likely existed at low frequency in the shared ancestral culture (indeed, we can detect one of the common polymorphisms, D106G, in individual clones at the Early time point, indicating that this mutation was heteroplasmic in cells of the founding population). Killer phenotypes are consistent across populations that fixed a particular ancestral polymorphism (Table S1).

In addition to the three ancestral polymorphisms, we detect 34 putative *de novo* point mutations that arose during the evolution of individual populations (Table S1). Mutations are localized to the K1 coding sequence, scattered across the four encoded subunits, and skewed towards missense mutations relative to nonsense or frameshift (Fig. 3B). Fourteen of the seventy-eight identified mutations are predicted to fall at or near sites of protease cleavage or post-translational modification necessary for toxin maturation. Overall, however, the K1 coding sequence appears to be under balancing selection (dN/dS=0.90), indicating that certain amino acid substitutions (e.g. those that eliminate immunity but retain killing ability) are not tolerated. In addition, substitutions are extremely biased toward transitions over transversions (Table S2, R=6.4, χ^2^=44.2, df=1, *P*<0.0001), a bias that is also present in other laboratory-derived M1 variants (R=4.1) (Suzuki et al. 2015) and natural variation of the helper L-A virus (R=3.0) (Diamond et al. 1989, Icho and Wickner 1989). The transition:transversion bias appears specific to viral genomes as the ratio is much lower within evolved nuclear genomes (R=0.8), especially in genes inferred to be under selection (R=0.5), suggesting a mutational bias of the viral RNA-dependent RNA polymerase (Lang et al. 2013, Fisher et al. 2018, Marad et al. 2018).

Though point mutations are the most common form of evolved variation, we also detected two viral genomes in which large portions of the K1 ORF are deleted (Fig. 3B). Despite the loss of the majority of the K1 coding sequence, the deletion mutants maintain cis signals for replication and packaging (Ribas and Wickner 1992, Ribas et al. 1994). Notably, the two populations that possess these deletion mutants also possess full-length viral variants. The deletion mutants we observe are similar to the ScV-S defective interfering particles that have been shown to outcompete full-length virus presumably due to their decreased replication time (Kane et al. 1979, Ridley and Wickner 1983, Esteban and Wickner 1988).

### Host/virus co-evolutionary dynamics are complex and operate over multiple scales

To compare the dynamics of viral genome evolution, nuclear genome evolution, and phenotypic evolution we performed time-course sequencing of viral genomes from three yeast populations that lost killing ability and for which we have whole-population, whole-genome, time-course sequencing data for the nuclear genome (Lang et al. 2013). As with the evolutionary dynamics of the host genome, the dynamics of viral genome evolution feature clonal interference (competition between mutant genotypes), genetic hitchhiking (an increase in frequency of an allele due to genetic linkage to a beneficial mutation), and sequential sweeps (Fig. 4, Fig. S6). Interestingly, viral sweeps often coincide with nuclear sweeps.

**Fig. 4.**
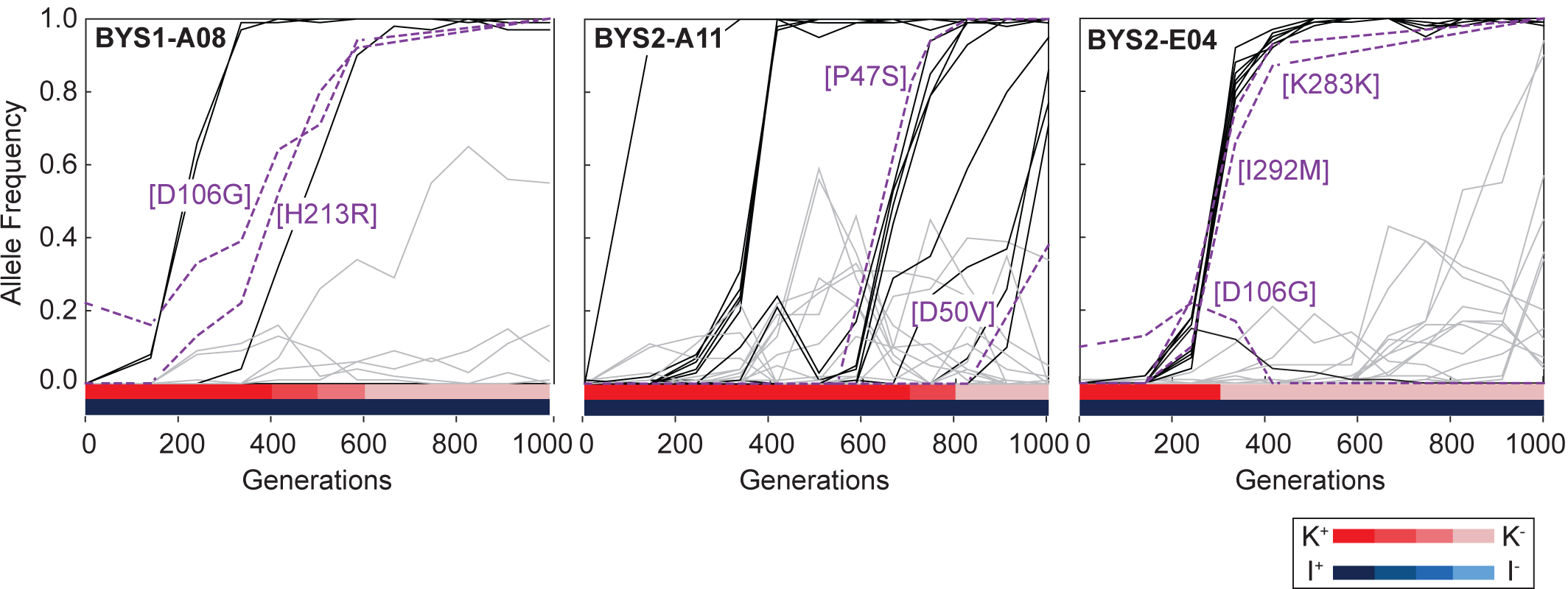
Viral dynamics mimic nuclear dynamics. Killer phenotype of evolved populations is indicated by color according to the key. Nuclear dynamics (reported previously Lang et al 2014) are represented as solid lines. Nuclear mutations that sweep before or during the loss of killing ability are indicated by black lines. All other mutations are indicated by gray lines. Viral mutations are indicated by purple dashed lines and labeled by amino acid change.

Since the coinciding nuclear sweeps often contain known driver mutations, it is possible that the viral variants themselves are not driving adaptation but instead hitchhiking on the back of beneficial nuclear mutations. This is consistent with the observation that the introduction of the viral variant from the Intermediate clone did not affect the fitness of the Early clone (Fig. 1B) To determine if the loss of killer phenotype is caused solely by mutations in the killer virus, we transferred the ancestral virus and five evolved viral variants to a virus-cured ancestor via cytoduction. Of the five viral variants one exhibited weak killing ability and full immunity (D253N), three exhibited no killing ability and full immunity (P47S, D106G, I292M), and one exhibited neither killing ability nor immunity (−1 frameshift). For each viral variant, the killer phenotype of the cytoductant matched the killer phenotype of the population of origin demonstrating that viral mutations are sufficient to explain changes in killer phenotype (Fig. S7). To determine if any viral variants affect host fitness, we competed all five cytoductants against the ancestor or a virus-cured ancestor in a pairwise manner. In competitions in which both competitors shared either killing ability or immunity, no viral variants impacted host fitness; therefore, production of active toxin or maintenance of the virus has no detectable fitness cost to the host (Fig. 5A). In contrast, an incompatibility in the killer phenotype – produced by the simultaneous loss of both killing ability and immunity – results in frequency dependent fitness interactions between competing populations. Collectively, these pairwise competitions show that a stepwise degradation of the killer virus is a neutral process and no evidence that host fitness is driving the loss of killer phenotype. These findings support previous theoretical and empirical studies (Pieczynska et al. 2016, Pieczynska et al. 2017) that claim that mycoviruses and their hosts have co-evolved to minimize cost.

**Fig. 5.**
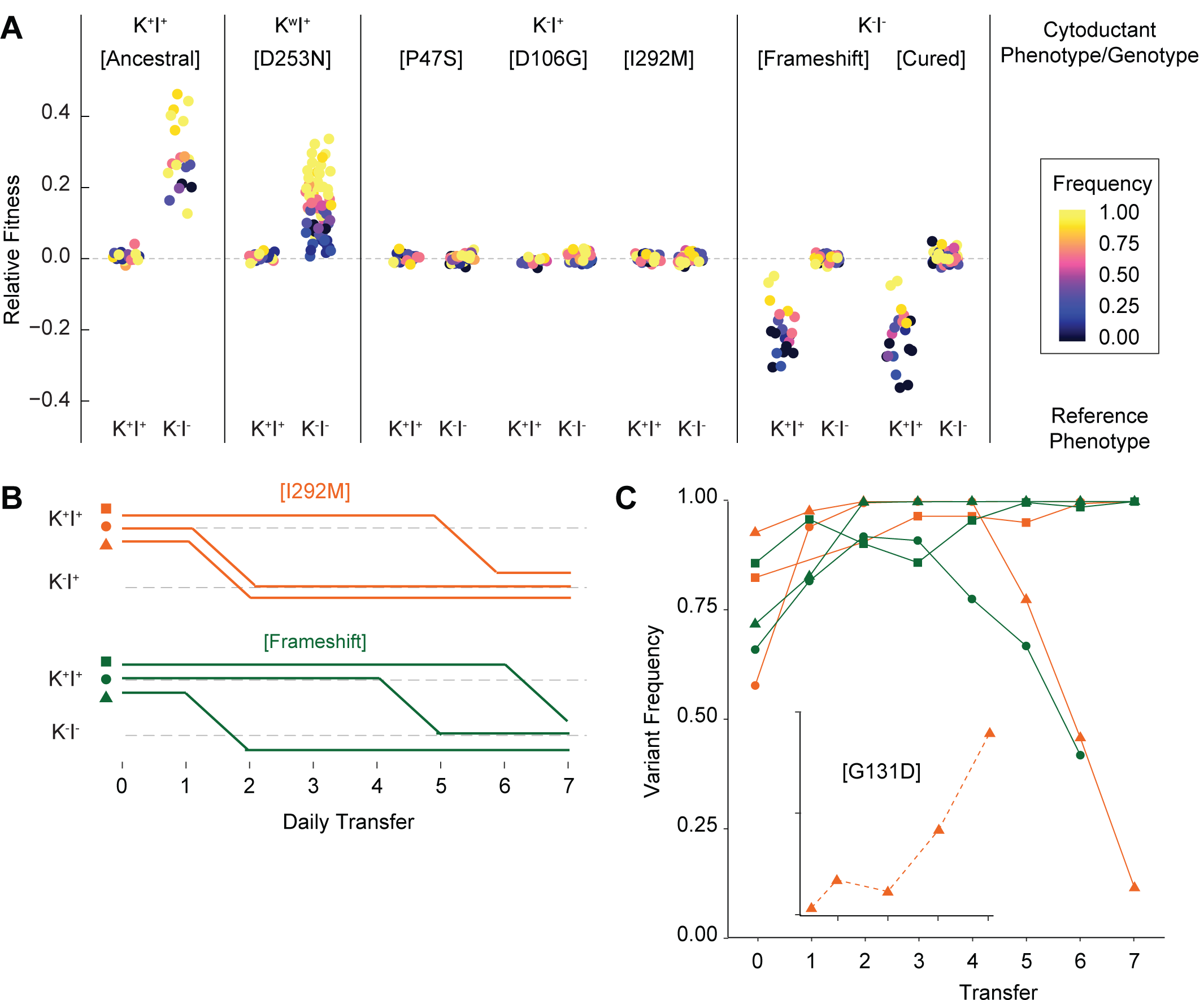
Viral evolution is driven by selection for an intracellular competitive advantage. A) Relative fitness of viral variants in pairwise competition with the ancestor (K^+^I^+^) and virus-cured ancestor (K^-^I^-^). Killer phenotype and identity of viral variant labeled above (K^w^ indicates weak killing ability). Killer phenotype of the ancestral competitor labeled below. Starting frequency indicated by color. B) Change to killer phenotype during intracellular competitions between viral variants (by color) and ancestral virus. Replicate lines indicated by symbol. C) Variant frequency during intracellular competitions. Colors and symbols consistent with panel B. Inset: frequency of the de novo G131D viral variant.

### Success of evolved viral variants is due to an intracellular fitness advantage

Based on the lack of a measurable effect of viral mutations on host fitness, we hypothesized that the evolved viral variants may have a selective advantage within the viral population of individual yeast cells. A within-cell advantage has been invoked to explain the invasion of internal deletion variants (e.g. ScV-S (Kane et al. 1979)) but has not been extended to point mutations. To test evolved viral variants for a within-cell fitness advantage, we generated a heteroplasmic diploid strain by mating the ancestor (with wildtype virus) with a haploid cytoductant containing either the I292M (K^-^I^+^) or -1 frameshift (K^-^I^-^) viral variant. The heteroplasmic diploids were propagated for seven single-cell bottlenecks every 48 hours to minimize among-cell selection. At each bottleneck, we assayed the yeast cells for killer phenotypes and we quantified the ratio of the intracellular viral variants by RT-PCR and sequencing. We find that killing ability was lost from all lines, suggesting that the evolved viral variants outcompeted the ancestral variant (Fig. 5B). Sequencing confirmed that the derived viral variant fixed in most lines (Fig. 5C). In some lines, however, the derived viral variant increased initially before decreasing late. Further investigation into one of these lines revealed that the decrease in frequency of the viral variant corresponded to the sweep of a *de novo* G131D variant (Fig. 5C, inset). Viral variants therefore appear to constantly arise, and the evolutionary success of the observed variants results from their selective advantage over viral competitors within the context of an individual cell. We speculate that an intracellular competition between newly arising viral variants also explains the loss of immunity from populations that previously lost killing ability (Fig. S5), given the relaxed selection for the maintenance of functional immunity in those populations.

## Discussion

We examined phenotypic and sequence co-evolution of an intracellular double-stranded RNA virus and the host nuclear genome over the course of 1,000 generations of experimental evolution. We observe complex dynamics including genetic hitchhiking and clonal interference in the host populations as well as the intracellular viral populations. Phenotypic and genotypic changes including the loss of killing ability, mutations in the host-encoded cell-wall biosynthesis genes, and the virally encoded toxin genes occur repeatedly across replicate populations. The loss of killer-associated phenotypes—killing ability and immunity to the killer toxin—leads to three phenomena with implications for adaptive evolution: positive frequency-dependent selection, multilevel selection, and nontransitivity.

Frequency-dependent selection can be either negative, where rare genotypes are favored, or positive, where rare genotypes are disfavored. Of the two, negative frequency-dependent selection is more commonly observed in experimental evolution, arising, for example, from cross-feeding between acetate producers and consumers (Helling et al. 1987, Turner et al. 1996, Spencer et al. 2008, Plucain et al. 2014) and spatial structuring in static environments (Rainey and Travisano 1998, Frenkel et al. 2015). Positive frequency-dependent selection, in contrast, is not typically observed in experimental evolution. By definition, a new positive frequency-dependent mutation must invade an established population at a time when its fitness is at its minimum. Even in situations in which positive frequency-dependent selection is likely to occur, such as the evolution of cooperative group behaviors and interference competition (Chao and Levin 1981), a mutation may be unfavorable at the time it arises. A crowded, structured environment provides an opportunity for allelopathies to offer a local advantage. Here we describe an alternative mechanism for the success of positive frequency-dependent mutations through multilevel selection of the host genome and a toxin-encoding intracellular virus. The likelihood of such a scenario occurring is aided by the large population size of the extrachromosomal element: each of the ∼10^5^ cells that comprise each yeast population contains ∼10^2^ viral particles (Bostian et al. 1983, Ridley and Wickner 1983).

Nontransitivity in our experimental system is due, in part, to interference competition: the production of a killer toxin that kills non-killer-containing cells. Interference competition can drive ecological nontransitivity (Kerr et al. 2002, Kirkup and Riley 2004), suggesting that similar mechanisms may underlie both ecological and genealogical nontransitivity. The adaptive evolution of genealogical nontransitivity in our system does not follow the canonical model of a co-evolutionary arms race where the host evolves mechanisms to prevent the selfish replication of the virus and the virus evolves to circumvent the host’s defenses (Daugherty and Malik 2012, Rowley 2017). Rather, mutations that fix in the viral and yeast populations do so because they provide a direct fitness advantage in their respective populations. Nontransitivity arises through the combined effect of beneficial mutations in the host genome (which improves the relative fitness within the yeast population, regardless of the presence or absence of the killer virus) and the adaptive loss of killing ability and degeneration of the intracellular virus (which provides an intracellular fitness advantage to the virus). The end result is a high-fitness yeast genotype (relative to the ancestral yeast genotype) that contains degenerate viruses, rendering their hosts susceptible to the virally-encoded toxin.

Though we did not find an impact of nuclear mutations on killer-associated phenotypes, we do observe a statistical enrichment of mutations in genes involved in β-glucan biosynthesis and in genes that when deleted confer a high level of resistance to the killer toxin. Nearly all mutations in these toxin-resistance genes are nonsynonymous (18 nonsense/frameshift, 21 missense, 1 synonymous), indicating a strong signature of positive selection. This suggests that the nuclear genome adapting in response to the presence of the killer toxin, however, the effect of these mutations may be beyond the resolution of our fitness assay.

Among the viral variants, we identified were two unique ∼1 kb deletions; remnants of the killer virus that retain little more than the *cis*-acting elements necessary for viral replication and packaging. These defective interfering particles are thought to outcompete full-length virus due to their decreased replication time (Kane et al. 1979, Ridley and Wickner 1983, Esteban and Wickner 1988). Defective interfering particles are common to RNA viruses (Holland et al. 1982). Though there are several different killer viruses in yeast (e.g. K1, K2, K28, Klus), each arose independently and has a distinct mechanism of action (Rodríguez-Cousiño et al. 2017). Nontransitive interactions may therefore arise frequently through cycles of gains and losses of toxin production and toxin immunity in lineages that contain RNA viruses.

Reports of nontransitivity arising along evolutionary lines of descent are rare (de Visser and Lenski 2002, Beaumont et al. 2009). The first (and most widely cited) report of nontransitivity along a direct line of descent occurred during yeast adaptation in glucose-limited chemostats (Paquin and Adams 1983). This experiment was correctly interpreted under the assumption—generally accepted at the time— that large asexual populations evolved by clonal replacement. This strong selection/weak mutation model, however, is now known to be an oversimplification for large asexual populations, where multiple beneficial mutations arise and spread simultaneously through the population (Gerrish and Lenski 1998, Kvitek and Sherlock 2013, Lang et al. 2013). In addition, the duration of the Paquin and Adams experiment was too short for the number of reported selective sweeps to have occurred (four in 245 generations and six in 305 generations, for haploids and diploids, respectively). The strongest known beneficial mutations in glucose-limited chemostats, hexose transporter amplifications, provide a fitness advantage of ∼30% (Gresham et al. 2008, Kvitek and Sherlock 2011) and would require a minimum of ∼150 generations to fix in a population size of 4 × 10^9^ (Otto and Whitlock 1997). We contend that Paquin and Adams observed nontransitive interactions among contemporaneous lineages—ecological nontransitivity—rather than nontransitivity among genealogical descendants. Apart from the present study, there are no other examples of nontransitivity arising along a line descent, but numerous examples of nontransitive interactions among contemporaneous lineages (Sinervo and Lively 1996, Kerr et al. 2002, Kirkup and Riley 2004, Károlyi et al. 2005, Laird and Schamp 2006, Reichenbach et al. 2007, Precoda et al. 2017, Menezes et al. 2019).

Here we present a mechanistic case study on the evolution of nontransitivity along a direct line of genealogical descent, and we determine the specific genetic events that lead to nontransitivity in one focal population. Our results show that the continuous action of selection can give rise to genotypes that are less fit compared to a distant ancestor. We show that nontransitive interactions can arise quickly due to multilevel selection in a host/virus system. In the context of this experiment multi-level selection is common—most yeast populations fix nuclear and viral variants by Generation 1,000. Overall, our results demonstrate that adaptive evolution is capable of giving rise to nontransitive fitness interactions along an evolutionary lineage, even under simple laboratory conditions.

## Supporting information

Combined Supplemental Materials

## Acknowledgments

We thank Reed Wickner, Amber Rice, and members of the Lang Lab for their comments on the manuscript. This work was supported by the NIH grant 1R01GM127420. Illumina data of viral competitions and evolved nuclear genomes are accessible under the BioProject ID PRJNA553562 and PRJNA205542, respectively. Conceptualization and writing were performed by S.W.B. and G.I.L. Investigation was performed by S.W.B. and A.B.R.

## Methods

### Growth Conditions and Strain Construction

Unless specified otherwise, yeast strains were propagated at 30°C in YPD + A&T (yeast extract, peptone, dextrose plus 100 μg/ml ampicillin and 25 μg/ml tetracycline to prevent bacterial contamination).

The ancestor and evolved populations were described previously (Lang et al. 2011). Early, Intermediate, and Late clones were isolated by resurrecting population BYS1-D08 at the Generation 0, 335, and 1,000, respectively. These specific timeponts were selected to coincide with the completion of a selective sweep (Lang et al. 2013), when the population is expected to be near clonal. For each timepoint we isolated multiple clones from a YPD plate and assayed each one to verify that the killer phenotype was uniform.

The ancestral strain was cured of the M1 and LA viruses by streaking to single colonies on YPD agar and confirmed by halo assay, PCR, and sequencing. We integrated a constitutively-expressed fluorescent reporter (pACT1-ymCitrine) at the *CAN1* locus in the cured ancestral strain as well as the Intermediate (Generation 335) and Late (Generation 1,000) clones.

Karyogamy mutants were constructed by introducing the *kar1Δ15* allele by two-step gene replacement in the cured a *MAT*α version of the ancestor (Georgieva and Rothstein 2002). The *kar1Δ15*-containing plasmid pMR1593 (Mark Rose, Georgetown University) was linearized with BglII prior to transformation and selection on -Ura. Mitotic excision of the integrated plasmid was selected for plating on 5-fluorotic acid (5-FOA). Then we perform replaced NatMX with KanMX to enable selection for recipients during viral transfer.

### Fitness Assays

Competitive fitness assays were performed as described previously (Lang et al. 2011, Lang et al. 2013). To investigate frequency dependence, competitors were mixed at various ratios at the initiation of the experiment. Competitions were performed for 50 generations under conditions identical to the evolution experiment (Lang et al. 2011). Every 10 generations, competitions were diluted 1:1,000 in fresh media and an aliquot was sampled by BD FACS Canto II flow cytometer. Flow cytometry data was analyzed using FlowJo 10.3. Relative fitness was calculated as the slope of the change in the natural log ratio between the experimental and reference strain. To detect frequency-dependent selection, each 10-generation interval was analyzed independently to calculate starting frequency and fitness.

### Halo Assay

Killer phenotype was measured using a high-throughput version of the standard halo assay (Crabtree et al. 2019) and a liquid handler (Biomek FX). Assays were performed using YPD agar that had been buffered to pH 4.5 (citrate-phosphate buffer), dyed with methylene blue (0.003%), and poured into a 1-well rectangular cell culture plate.

Killing ability was assayed against a sensitive tester strain (yGIL1063) that was isolated from a separate evolution experiment initiated from the same ancestor. The sensitive tester was grown to saturation, diluted 1:10, and spread (150 µL) evenly on the buffered agar. Query strains were grown to saturation, concentrated 5x, and spotted (2 µL) on top of the absorbed lawn (Fig. S2, left).

Immunity was assayed against the ancestral strain (yGIL432). Query strains were grown to saturation, diluted 1:32, and spotted (10 µL) on the buffered agar. The killer tester was grown to saturation, concentrated 5x, and spotted (2 µL) on top of the absorbed query strain (Fig. S2, right). Plates were incubated at room temperature for 2-3 days before assessment. Killer phenotype was scored according to the scale in shown in Fig. 2.

### Viral RNA Isolation, cDNA Synthesis, PCR

Nucleic acids were isolated by phenol-chloroform extraction and precipitated in ethanol. Isolated RNA was reverse-transcribed into cDNA using ProtoScript II First Strand cDNA Synthesis Kit (NEB) with either the enclosed Random Primer Mix or the M1-specific oligo M1_R3 (Table S3).

### Sanger Sequencing and Bioinformatics Analyses

PCR was performed on cDNA using Q5 High-Fidelity Polymerase (NEB). The K1 ORF was amplified using primers M1_F1 or M1_F5 and M1_R6 (Table S3). The M1 region downstream of the polyA stretch was amplified using M1_F7 and M1_R3. The LA virus was amplified using LA_F2 and LA_R2, LA_F2 and LA_R3, or LA_F3 and LA_R6. PCR products were Sanger sequenced by Genscript.

Mutations were identified and peak height quantified using 4Peaks (nucleobytes). For intracellular competitions, mutation frequency was quantified by both Sanger and Illumina sequencing (see below), with both methods producing nearly identical results (Fig. S8).

The Sanger sequencing data was aligned to publicly-available M1 and LA references (GenBank Accession Numbers U78817 and J04692, respectively) using ApE (A plasmid Editor). The ancestral M1 and LA viruses differed from the references at 7 sites (including 3 K1 missense mutations) and 19 sites, respectively (Fig. S3).

### Viral Transfer

Viruses were transferred to *MAT***a** strains using the *MAT*α karyogamy mutant as an intermediate. Viral donors (*MAT***a**, *ura3*, NatMX) were first transformed with the pRS426 (*URA3*, 2µ ORI) for future indication of viral transfer. Cytoduction was performed by mixing a viral donor with the karyogamy mutant recipient (*MAT*α, *ura3*, KanMX) at a 5:1 ratio on solid media. After a 6 hr incubation at 30°C, the cells were plated on media containing G418 to select for cells with the recipient nuclei. Recipients that grew on -Ura (indicator of cytoplasmic mixing) and failed to grow on ClonNat (absence of donor nuclei) then served as donors for the next cytoduction. These karyogamy mutant donors (*MAT*α, *URA3*, KanMX) were mixed with the selected recipient (*MAT***a**, *ura3*, NatMX) at a 5:1 ratio on solid media. After a 6 hr incubation at 30°C, the cells were plated on media containing ClonNat to select for cells with recipient nuclei. Recipients that grew on -Ura (indicator of cytoplasmic mixing) and failed to grow on ClonNat (absence of the donor nucleus) were then cured of the indicator plasmid by selection on 5-FOA. Killer phenotype was confirmed by halo assays and the presence of the viral variants in the recipient was verified by Sanger sequencing.

### Illumina Sequencing and Bioinformatics Analyses

Multiplexed libraries were prepared using a two-step PCR. First, cDNA was amplified by Q5 High-Fidelity Polymerase (NEB) for 10 cycles using primers I292M_read1 and I292M_read2 or frameshift_read1 and frameshift_read2 (Table S3) to incorporate a random 8 bp barcode and sequencing primer binding sites. The resulting amplicons were further amplified by Q5 PCR for 15 cycles using primers i5_adapter and i7_adapter to incorporate the sequencing adaptors and indices. Libraries were sequenced on a NovaSeq 6000 (Illumina) at the Genomics Core Facility at Princeton University.

Raw FASTQ files were demultiplexed using a dual-index barcode splitter (https://bitbucket.org/princeton_genomics/barcode_splitter) and trimmed using Trimmomatic (Bolger et al. 2014) with default settings for paired-end reads. Mutation frequencies were determined by counting the number of reads that contain the ancestral or evolved allele (mutation flanked by five nucleotides).

### Intracellular Competitions

Within-cell viral competitions were performed by propagating a heteroplasmic diploid and monitoring killer phenotype and viral variant frequency. Diploids were generated by crossing the ancestor with a cytoductant harboring either the I292M or -1 frameshift viral variant. For each viral variant, three diploid lines (each initiated by a unique mating event) were passaged every other day on buffered YPD media for a total of 7 single-cell bottlenecks to minimize among-cell selection. A portion of each transferred colony was cryopreserved in 15% glycerol. Cryosamples were revived, assayed for killer phenotype, and harvested for RNA. Following RT-PCR, samples were sent for Sanger sequencing and Illumina sequencing. Variant frequency deviated from the expected frequency of 0.5 at the initial timepoint, presumably due to an unavoidable delay between the formation of the heteroplasmic diploid and initiation of the intracellular competition from a single colony. Alternatively, viral copy number may vary between donor and recipient cells.

