## Supplementary material for "Adaptive evolution of nontransitive fitness in yeast": Combined Supplemental Materials

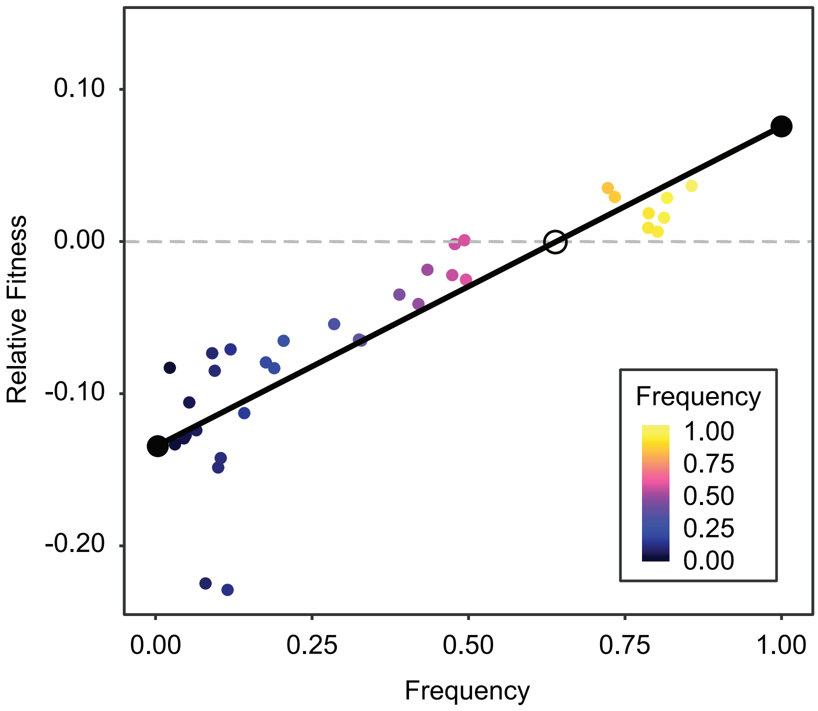


**Fig. S1.** Positive frequency dependent interaction along an evolutionary lineage. Fitness of Late clone relative to Early clone, as a function of frequency. Stable fixed points indicated by closed black circles and unstable fixed point indicated by open black circle.

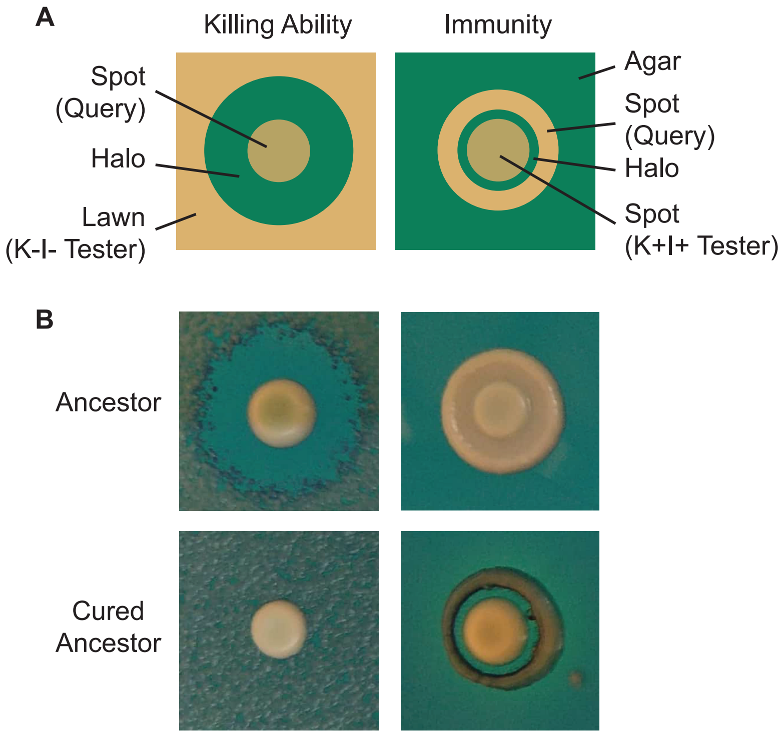


**Fig. S2.** Visualization of killer phenotype by halo assay. A) Schematic of killer phenotypic assays. To assay killing ability, a tester (sensitive) strain is spread as a lawn, followed by a query strain spotted as a concentrated culture. After incubation, the production of a zone of clearing indicates that the query strain possesses killing ability. To assay sensitivity, a query strain is plated as a dilute spot, followed by a tester (killer) strain spotted as a concentrated culture. After incubation, the production of a zone of clearing indicates that the query strain possesses killing ability. B) Halo assays demonstrate that the ancestor of the evolution experiment exhibits killing ability and immunity while the cured ancestor lacks killing ability and immunity.

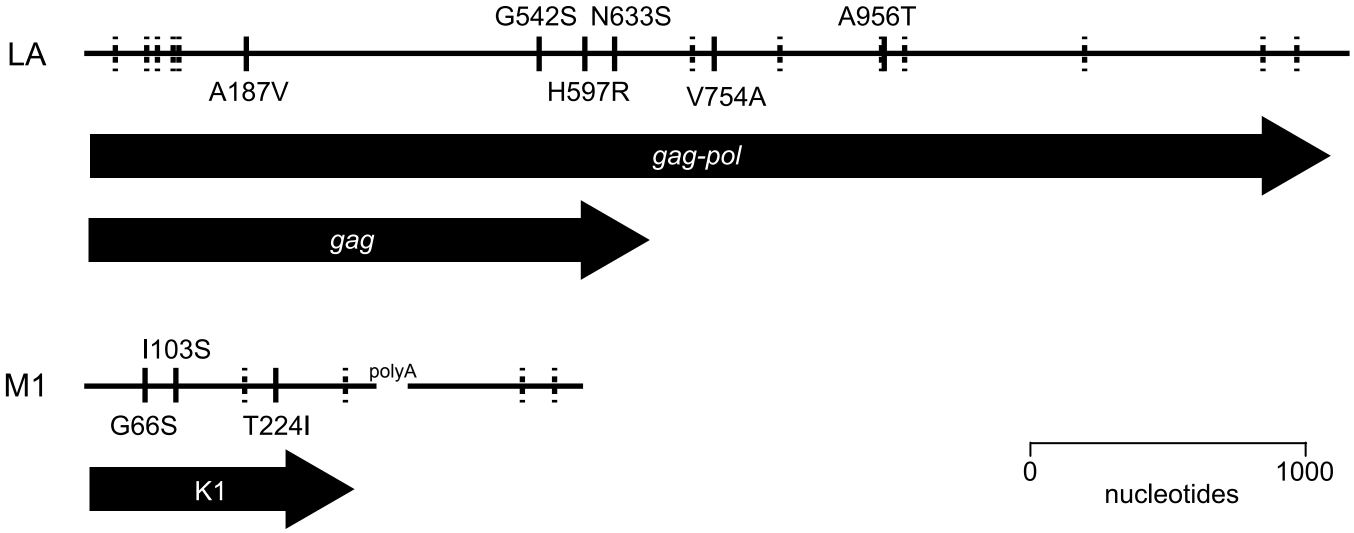


**Fig. S3.** Sequence divergence of ancestral viruses. The viruses of our ancestral yeast strain diverged from previously published LA and M1 genomes by 19 nucleotides and 7 nucleotides, respectively. Solid lines represent nonsynonymous polymorphisms, labeled by amino acid substitution. Dashed lines represent synonymous/intergenic polymorphisms.

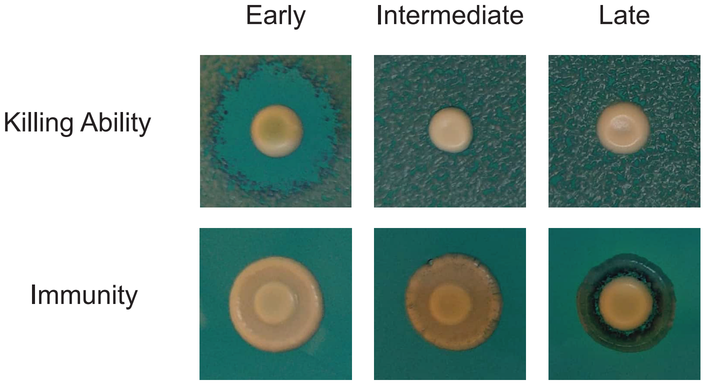


**Fig. S4.** Stepwise deterioration of killer phenotype in evolved clones. The killer phenotypes of Early, Intermediate, and Late clones from population BYS1-D08 were determined by halo assay.

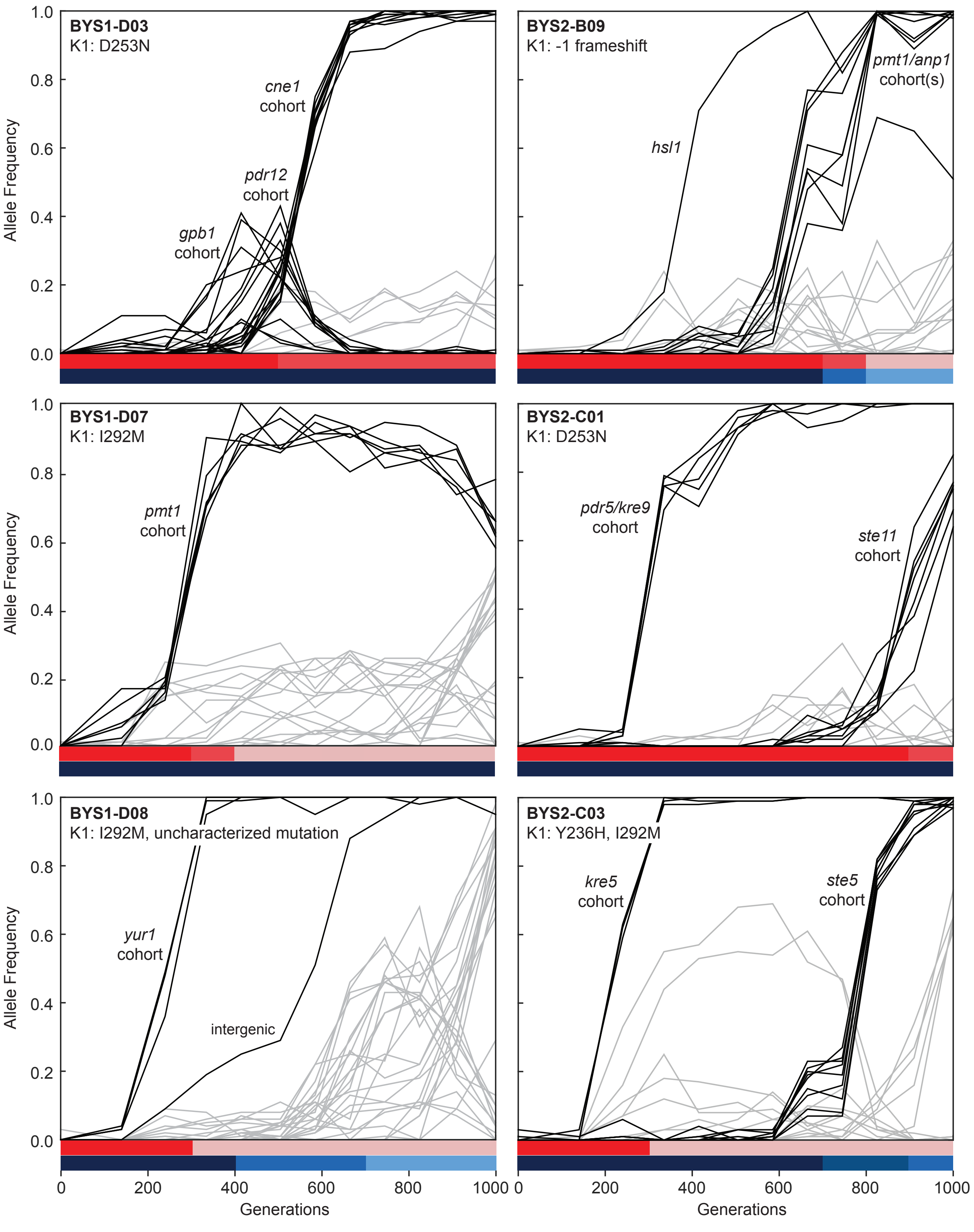


**Fig. S5.** Evolutionary dynamics of nuclear genotypes and killer phenotypes over time. The K1 mutations detected in each population at Generation 1,000 are indicated in the top-left of the plot. Trajectories of nuclear mutations were obtained from Lang et al. 2013. Black lines indicate nuclear mutations that swept up to and including the period of killer phenotypic change (all others nuclear mutations are gray). Mutational cohorts are labeled according to their putative driver or putative toxin resistance mutation. Killing ability and immunity are indicated in bar graph (bottom) by shades of red and blue, respectively.

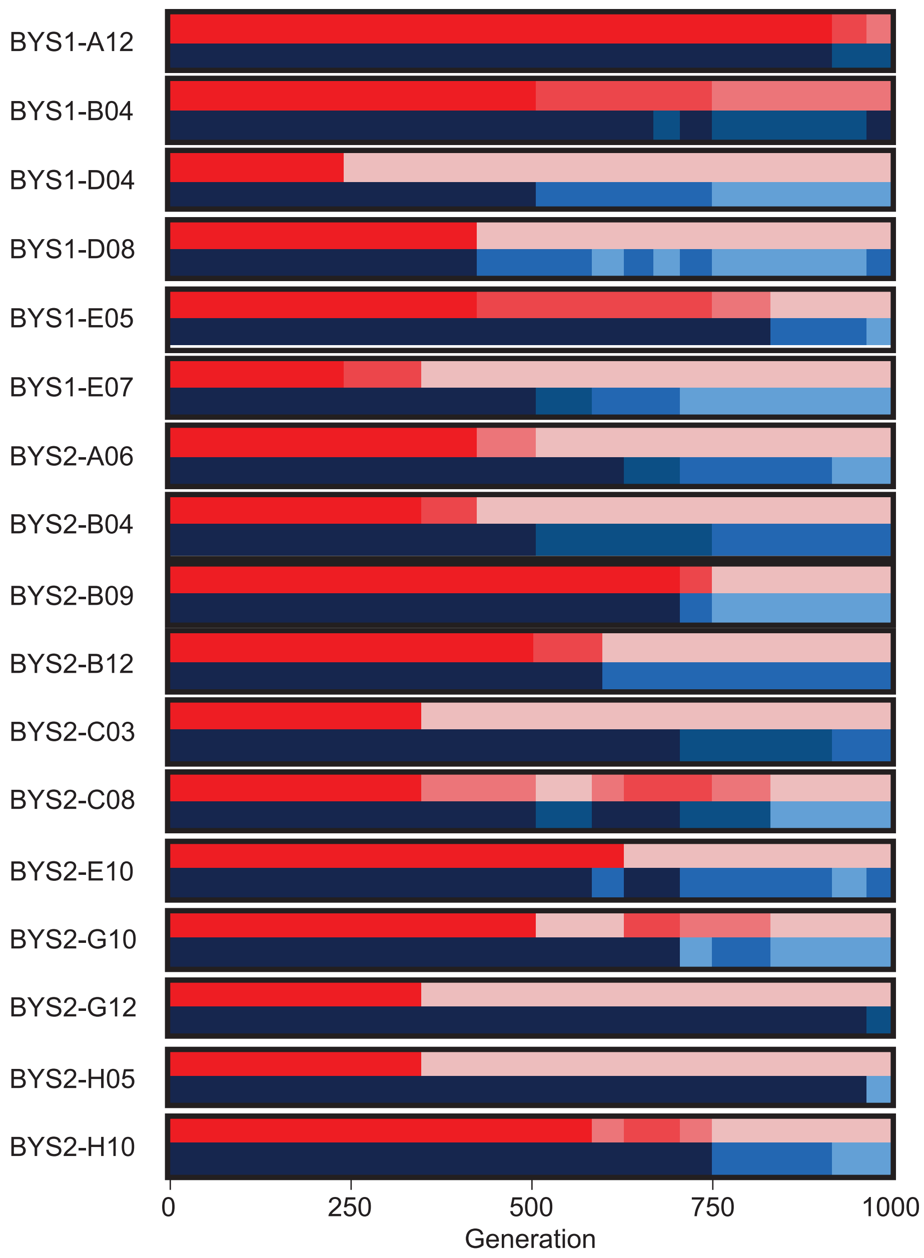


**Fig. S6.** Killer phenotypes of the 17 populations that develop sensitivity to the K1 toxin. Killer phenotype is shown according to scale in Fig. 2. For each population, killing ability is shown in shades of red (top) and immunity in shades of blue (bottom).

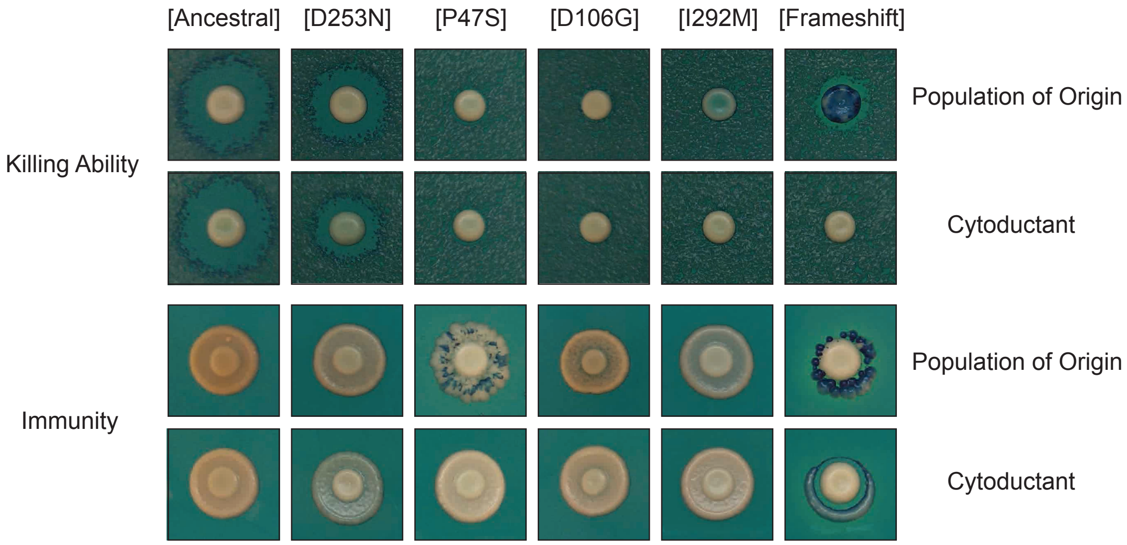


**Fig. S7.** Cytoductants exhibit the same killer phenotype as the population of origin. Viral variants were transferred from evolved populations to a cured ancestor. Halo assays demonstrate that killer phenotypes were consistent between donor and recipient strains. Viruses were obtained from the following evolved populations at Generation 1,000: BYS1-A03 (D253N), RMB1-A02 (P47S), BYB1-H06 (D106G), BYS1-A05 (I292M), BYS2-B09 (frameshift). Populations RMB1-A02 and BYS2-B09 appear mixed given the observed speckling pattern.

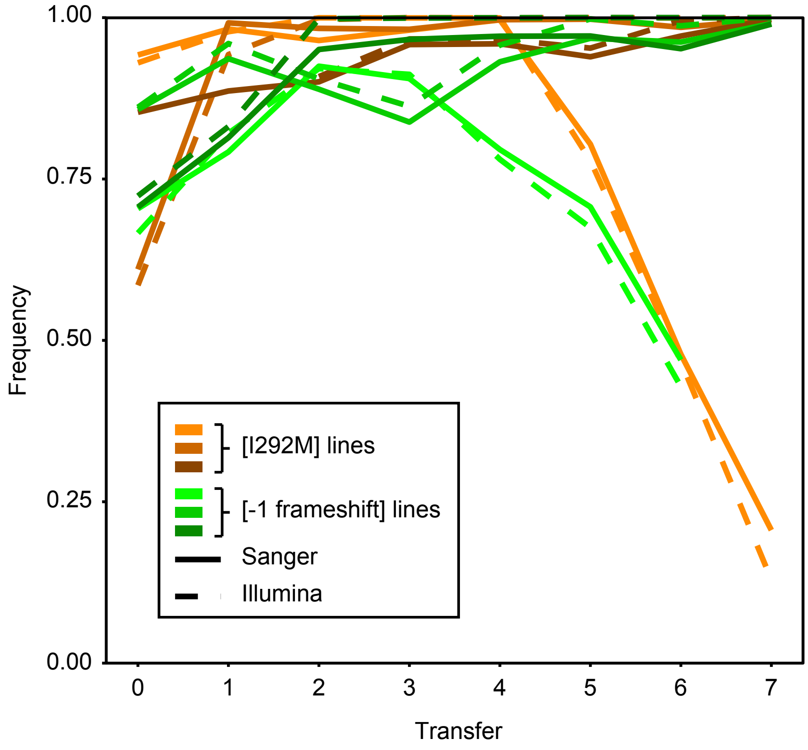


**Fig. S8.** Consensus between Sanger and Illumina sequencing in reporting mutation frequency. Intracellular competitions were tracked over time by both Sanger and Illumina sequencing.

**Change Killer Phenotype**

| Population | Killing Ability | Immunity | K1 Mutations (subunit) |
|---|---|---|---|
| BYS1-D04 | 0.00 | 0.00 | D106G (α), Y185N (γ) |
| BYS1-E05 | 0.00 | 0.00 | I243I (β), D253N (β) |
| BYS1-E07 | 0.00 | 0.00 | M1, LA not detected |
| BYS2-A06 | 0.00 | 0.00 | G289S (β) |
| BYS2-B09 | 0.00 | 0.00 | -1 bp frameshift (α) |
| BYS2-C08 | 0.00 | 0.00 | D253N (β) |
| BYS2-G10 | 0.00 | 0.00 | I292M (β) |
| BYS2-H10 | 0.00 | 0.00 | I292M (β) |
| BYS1-D08 | 0.00 | 0.33 | I292M (β), uncharacterized mutation* |
| BYS2-B04 | 0.00 | 0.33 | Y236C (β) |
| BYS2-C03 | 0.00 | 0.33 | Y236H (β), I292M (β) |
| BYS2-E10 | 0.00 | 0.33 | D106G (α), K296E (β) |
| BYS1-A12 | 0.33 | 0.67 | L27P (δ), N90N (α) |
| BYS1-A02 | 0.00 | 1.00 | S13S (δ), D106G (α) |
| BYS1-A05 | 0.00 | 1.00 | I292M (β) |
| BYS1-A08 | 0.00 | 1.00 | D106G (α), H213R (γ) |
| BYS1-B02 | 0.00 | 1.00 | D106G (α) |
| BYS1-B03 | 0.00 | 1.00 | I292M (β) |
| BYS1-B06 | 0.00 | 1.00 | D168G (γ), I292M (β) |
| BYS1-C02 | 0.00 | 1.00 | I292M (β) |
| BYS1-C06 | 0.00 | 1.00 | I292M (β) |
| BYS1-C10 | 0.00 | 1.00 | S66S (α), D106G (α) |
| BYS1-D06 | 0.00 | 1.00 | D106G (α), Δ 878 bp ^ |
| BYS1-D07 | 0.00 | 1.00 | I292M (β) |
| BYS1-G07 | 0.00 | 1.00 | D106G (α) |
| BYS2-A01 | 0.00 | 1.00 | I292M (β) |
| BYS2-A04 | 0.00 | 1.00 | T36T (δ), Y236H (β), I292M (β) |
| BYS2-A11 | 0.00 | 1.00 | P47S (α), D50V (α) |
| BYS2-B11 | 0.00 | 1.00 | D106G (α) |
| BYS2-C05 | 0.00 | 1.00 | K244R (β), V256M (β) |
| BYS2-C09 | 0.00 | 1.00 | T195T (γ), I292M (β) |
| BYS2-E02 | 0.00 | 1.00 | D106G (α) |
| BYS2-E04 | 0.00 | 1.00 | K283K (β), I292M (β) |
| BYS2-E06 | 0.00 | 1.00 | D106G (α), F222L (γ) |
| BYS2-E07 | 0.00 | 1.00 | D106G (α) |
| BYS2-E11 | 0.00 | 1.00 | I292M (β), Δ1177 bp ^ |
| BYS1-B04 | 0.33 | 1.00 | D253N (β) |
| BYS1-B08 | 0.33 | 1.00 | R149H (γ) |
| BYS1-B10 | 0.33 | 1.00 | Y282C (β) |

| | | | |
|---|---|---|---|
| BYS1-D10 | 0.33 | 1.00 | W277R (β) |
| BYS1-E08 | 0.33 | 1.00 | Q184R (γ), N307D (β), D308N (β) |
| BYS2-A03 | 0.33 | 1.00 | D106G (α) |
| BYS2-F05 | 0.33 | 1.00 | D253N (β) |
| BYS2-G05 | 0.33 | 1.00 | Y219H (γ) |
| BYS1-A03 | 0.67 | 1.00 | D253N (β) |
| BYS1-A09 | 0.67 | 1.00 | N280D (β) |
| BYS1-C05 | 0.67 | 1.00 | D253N (β) |
| BYS1-D03 | 0.67 | 1.00 | D253N (β) |
| BYS1-E02 | 0.67 | 1.00 | T223I (γ) |
| BYS1-H07 | 0.67 | 1.00 | W291R (β) |
| BYS1-H11 | 0.67 | 1.00 | D253N (β) |
| BYS2-A10 | 0.67 | 1.00 | T64T (α), D253N (β) |
| BYS2-B08 | 0.67 | 1.00 | D253N (β) |
| BYS2-C01 | 0.67 | 1.00 | D253N (β) |
| BYS2-C12 | 0.67 | 1.00 | D253N (β) |
| BYS2-D02 | 0.67 | 1.00 | D253N (β) |
| BYS2-F02 | 0.67 | 1.00 | H214H (γ), D253N (β) |

**Retain Killer Phenotype**

| Population | Killing Ability | Immunity | K1 Mutations (subunit) |
|---|---|---|---|
| BYS1-A04 | 1.00 | 1.00 | |
| BYS1-A07 | 1.00 | 1.00 | |
| BYS1-F05 | 1.00 | 1.00 | |
| BYS1-G01 | 1.00 | 1.00 | |
| BYS1-G02 | 1.00 | 1.00 | |
| BYS2-C06 | 1.00 | 1.00 | L269L (β) |
| BYS2-D06 | 1.00 | 1.00 | |
| BYS2-D07 | 1.00 | 1.00 | |
| BYS2-E01 | 1.00 | 1.00 | |
| BYS2-E03 | 1.00 | 1.00 | |

**Table S1.** Killer phenotype and K1 mutations in evolved yeast populations at Generation 1,000. A caret (^) indicates that a population is heteroplasmic for variants listed. An asterisk (*) indicates that the mutation results in loss of PCR primer binding sites thereby preventing further characterization.

| **Nucleotide Substitution** | **K1** (this study) | **K1** (*12*) | **Gag-Pol LA** (*13,14*) | **Nuclear** (*6,15,16*) | **Nuclear – Common Targets** (*6,15,16*) |
|---|---|---|---|---|---|
| C:G>U:A (Ts) | 10 (27%) | 17 (33%) | 17 (35%) | 1174 (31%) | 47 (23%) |
| A:U>G:C (Ts) | 22 (59%) | 24 (47%) | 19 (40%) | 558 (15%) | 24 (12%) |
| A:U>C:G (Tv) | 1 (3%) | 5 (10%) | 1 (2%) | 348 (9%) | 11 (5%) |
| A:U>U:A (Tv) | 4 (11%) | 0 (0%) | 6 (13%) | 368 (10%) | 20 (10%) |
| C:G>A:U (Tv) | 0 (0%) | 4 (8%) | 4 (8%) | 850 (22%) | 65 (32%) |
| C:G>G:C (Tv) | 0 (0%) | 1 (2%) | 1 (2%) | 530 (14%) | 35 (17%) |
| Transitions | 32 | 41 | 36 | 1732 | 71 |
| Transversions | 5 | 10 | 12 | 2096 | 131 |
| R | 6.4 | 4.1 | 3.0 | 0.8 | 0.5 |

**Table S2.** Mutational biases in viral and nuclear datasets.

| Name | Sequence (5' → 3') | Use |
|---|---|---|
| M1_F1 | TTGGCTATTACAGCGTGCCA | PCR (Sanger) |
| M1_F5 | ATGACGAAGCCAACCCAAGT | PCR (Sanger) |
| M1_F7 | CAGAAAAAGAGAGAACAGGAC | PCR (Sanger) |
| M1_R3 | TGCTGTTGCATTAAACCAGGC | cDNA synthesis, PCR (Sanger) |
| M1_R6 | ATAGCCCGGTGCTCTGTAGG | PCR (Sanger) |
| LA_F2 | ATCAGGTGATGCAGCGTTGA | PCR (Sanger) |
| LA_F3 | ACTCCCCATGCTAAGATTTGTT | PCR (Sanger) |
| LA_R2 | CGGCACCCTTACGGAGATAC | PCR (Sanger) |
| LA_R3 | GACCTGTAATGCCCGGAGTG | PCR (Sanger) |
| LA_R6 | AGTACTGAGCCCCAAGACCA | PCR (Sanger) |
| I292M_read1 | CGTCGGCAGCGTCAGATGTGTATAAGAGACAGNNNNNNNNCCATGGTGTCGGCTAATGGT | PCR (Illumina) |
| I292M_read2 | CGTGGGCTCGGAGATGTGTATAAGAGACAGAGGTCAGACACGATGCCCTA | PCR (Illumina) |
| frameshift_read1 | CGTCGGCAGCGTCAGATGTGTATAAGAGACAGNNNNNNNNCCCGTCTGCGACAGTAGAAA | PCR (Illumina) |
| frameshift_read2 | CGTGGGCTCGGAGATGTGTATAAGAGACAGTGTGTAAGAACTGCGTGGGT | PCR (Illumina) |
| i5_adapter | **AATGATACGGCGACCACCGAGATCTACAC**NNNNNNNNTCGTCGGCAGCGTCAGATG | PCR (Illumina) |
| i7_adapter | **CAAGCAGAAGACGGCATACGAGAT**NNNNNNNNGTCTCGTGGGCTCGGAGATGTG | PCR (Illumina) |

**Table S3.** Oligos used in this study. primer binding sites are underlined and adaptor sequences are bolded. Sequences that correspond to barcodes and indices are shown as "N".
